# Paradoxical Associations Between Familial Affective Responsiveness, Stress, and Amygdala Reactivity

**DOI:** 10.1101/169383

**Authors:** Madeline J. Farber, Adrienne L. Romer, M. Justin Kim, Annchen R. Knodt, Nourhan M. Elsayed, Douglas E. Williamson, Ahmad R. Hariri

## Abstract

Studies of early life extremes such as trauma, abuse, and neglect highlight the critical importance of quality caregiving in the development of brain circuits supporting emotional behavior and mental health. The impact of normative variability in caregiving on such biobehavioral processes, however, is poorly understood. Here, we provide initial evidence that even subtle variability in normative caregiving shapes threat-related brain function and, potentially, associated psychopathology in adolescence. Specifically, we report that greater familial affective responsiveness is associated with heightened amygdala reactivity to interpersonal threat, particularly in adolescents having experienced relatively low recent stress. These findings extend the literature on the effects of caregiving extremes on brain function to subtle, normative variability, but suggest that presumably protective factors may be associated with increased risk-related amygdala reactivity. We consider these paradoxical associations with regard to studies of basic associative threat learning and further consider their relevance for understanding potential effects of caregiving on psychological development.

## Introduction

A wealth of empirical research demonstrates that the quality of caregiving predicts later psychosocial outcomes, in part, by shaping the development of brain circuits supporting emotional behavior (Belsky & de Haan, 2011; Callaghan & Tottenham, 2015; Cicchetti & Curtis, 2015). However, much of this research has focused on caregiving extremes, such as trauma, abuse, and neglect (Gee et al., 2013; Malter Cohen et al., 2013; Herringa et al., 2013; McLaughlin et al., 2015; Pechtel et al., 2014; Sheridan et al, 2012; Tottenham et al., 2010; Tottenham, 2012). Such a focus ignores the potential impact of normative variability in caregiving on neurodevelopment and behavior. A rich literature demonstrates lasting positive effects of normative caregiving including secure attachment and positive parenting (i.e., high care, low control) on behavioral development (Bowlby, 1978; Bowlby & Zeanah, 1988; Canetti et al., 1997; Francis & Meaney, 1999; Overbeek et al., 2007). Despite this literature, comparatively little is known about the potential impact of normative variability in caregiving on the underlying brain circuitry implicated in both normal and abnormal behavior (Romund et al., 2016; Tan et al., 2014; Whittle et al., 2009).

Here, we seek to address this research gap by examining how normative variability in caregiving within the family affects individual differences in threat-related amygdala reactivity in a large cohort of adolescents aged 12-15 years. We focus on threat-related amygdala reactivity because of converging evidence that this specific aspect of brain function is not only affected by extremes of caregiving (Gee et al., 2013; Malter Cohen et al., 2013; Herringa et al., 2013; McLaughlin et al., 2015; Pechtel et al., 2014; Sheridan et al, 2012; Tottenham et al., 2010; Tottenham, 2012) but also represents a biomarker of future risk for stress-related psychopathology (Admon et al., 2013; Etkin & Wager, 2007; Schlund & Cataldo, 2010; Swartz et al., 2015). In contrast to the extant literature, which represents a nearly exclusive focus on the effects of maternal caregiving (Caldji et al., 2015; Francis & Meaney, 1999; Gee et al., 2013; Romund et al., 2016; Tan et al., 2014; Tottenham et al., 2012; Whittle et al., 2009), we examine holistic family functioning to help inform more general effects of the caregiving environment on behaviorally and clinically relevant brain function.

Our analyses centered on data available for a subset of participants having completed the Teen Alcohol Outcomes Study, originally developed to examine the impact of familial loading for depression on the risk to develop alcohol use disorders. Study participants were characterized as either “high risk” (having a first- and second-degree relative with major depressive disorder; MDD) or “low risk” (having no first-degree and minimal second-degree relatives (< 20%) with MDD) (Williamson et al., 2004). The broader caregiving environment was indexed through the lens of family functioning as measured by the general functioning and affective responsiveness subscales of the Family Assessment Device (FAD). Amygdala reactivity to explicit, interpersonal and implicit, environmental threat as communicated by angry and fearful facial expressions, respectively, was measured using task-based functional MRI.

We hypothesized that lower scores on the FAD affective responsiveness subscale, which reflect situationally and contextually appropriate expression and recognition of emotion, would be associated with decreased amygdala reactivity specifically to angry facial expressions as canonical examples of explicit interpersonal threat. Our hypothesis is based on the assumption that interpersonal conflict is less likely to be experienced when family members appropriately express emotions, and that this positive modeling of emotional experiences would thus be reflected in lower amygdala reactivity to signals of such conflict. As an extension of this primary hypothesis, we further explored the following related questions. Are significant associations with amygdala reactivity specific to affective responsiveness and not more general family functioning? Moreover, are significant associations independent of early life stress, contemporaneous symptoms of depression and anxiety, and broad familial risk for depression? And, lastly, are significant associations moderated by recent stressful life events?

## Results

### Participant Characteristics

Demographic and behavioral characteristics of our analysis sample are detailed in Table 1. High and low risk groups significantly differed in symptoms of depression [*t*(200.4)=2.405, *p*=0.019] and in early life stress [*t*(212.2)=2.228, *p*=0.027]. However, the high and low risk groups did not differ in FAD general functioning [*t*(229)=-1.656, *p*=0.099] or affective responsiveness [*t*(229)=- 0.382, *p*=0.703]. Groups also did not differ in anxiety symptoms [*t*(217.3)=1.148, *p*=0.252] or recent life stress [*t*(210.3)=1.234, *p=*0.219]. There were no significant differences between groups on possible neuroimaging confounds including task performance as measured by accuracy [*t*(225)=-0.230, *p=*0.818] and response time [*t*(219)=-0.552, *p*=-0.582], or in task-related movement as indexed by head displacement [*t*(230)=-0.609, *p*=0.543].

**Table 1.** Sample Characteristics.

|  | **High Risk**<br>**(*n*=120)** |  | **Low Risk**<br>**(*n*=112)** |  | **Group**<br>**Comparisons** |  |
| --- | --- | --- | --- | --- | --- | --- |
|  | **Mean** | **SD** | **Mean** | **SD** | ***t*** | ***p*** |
| **Age** | 13.61 | 1.01 | 13.56 | 0.94 | 0.416 | 0.678 |
| **FAD Affective Responsiveness** | 16.69 | 2.72 | 16.84 | 2.96 | 0.483 | 0.630 |
| **FAD General Functioning** | 36.79 | 4.53 | 37.85 | 5.11 | -0.795 | 0.427 |
| **MFQ** | 4.06 | 4.20 | 2.96 | 2.60 | 2.405 | 0.017* |
| **SCARED** | 16.64 | 11.46 | 15.12 | 8.51 | 1.148 | 0.252 |
| **CTQ** | 34.55 | 6.85 | 32.59 | 6.01 | 2.228 | 0.027* |
| **SLES** | 13.40 | 10.22 | 12.20 | 8.44 | 1.234 | 0.219 |
| **Mean Accuracy (%)** | 95.72 | 17.42 | 96.23 | 15.92 | -0.237 | 0.818 |
| **Mean Response Time (ms)** | 1288.61 | 240.85 | 1306.66 | 245.35 | -0.522 | 0.582 |
| **Mean Head Displacement (mm)** | 0.04 | 0.01 | 0.04 | 0.02 | -0.609 | 0.543 |

### Family Functioning and Amygdala Reactivity

Across all participants, there was robust bilateral amygdala reactivity to both angry and fearful facial expressions (Figure 1). Linear regression analyses using extracted BOLD parameter estimates from clusters exhibiting main effects of expression revealed a significant association between FAD affective responsiveness and amygdala reactivity to angry facial expressions. Contrary to our hypothesis, participants who reported better affective responsiveness (i.e., lower scores) exhibited significantly higher reactivity to angry facial expressions in the left [*F*(1, 221)=4.305, *p*=0.039] but not right [*F*(1, 221)=1.005, *p*=0.317] hemisphere (Figure 2). This association was independent of broader familial risk for depression, as there was no statistically significant interaction between affective responsiveness and risk group status on reactivity [*F*(1, 221)=0.012, *p*=0.912]. Additional analyses revealed that the association between affective responsiveness and amygdala reactivity to angry facial expressions was further independent of early life stress [*F*(1, 204)=5.630, *p*=0.019] as well as contemporaneous symptoms of anxiety and depression [*F*(1, 203)=5.741, *p*=0.017].

**Figure 1.**
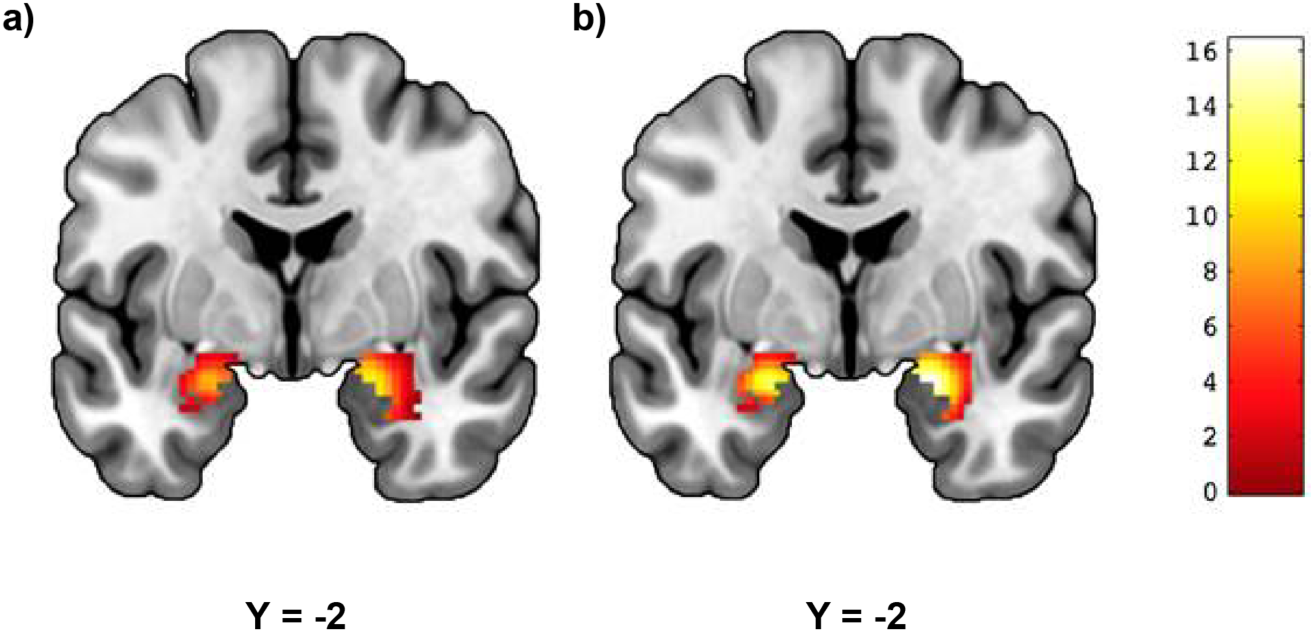
Threat-Related Amygdala Reactivity. Mean reactivity to (a) angry and (b) fearful facial expressions within anatomically defined amygdala regions of interest (*p*<0.05, FWE-corrected). (a) Left: cluster size = 169 voxels, peak voxel = -22, -2, -18; Right: cluster size = 226 voxels, peak voxel = 20, -4, -16; (b) Left: cluster size = 170 voxels, peak voxel = -22, -2, -18; Right: cluster size = 224 voxels, peak voxel = 22, -4, -18. Color bar represents t-scores.

**Figure 2.**
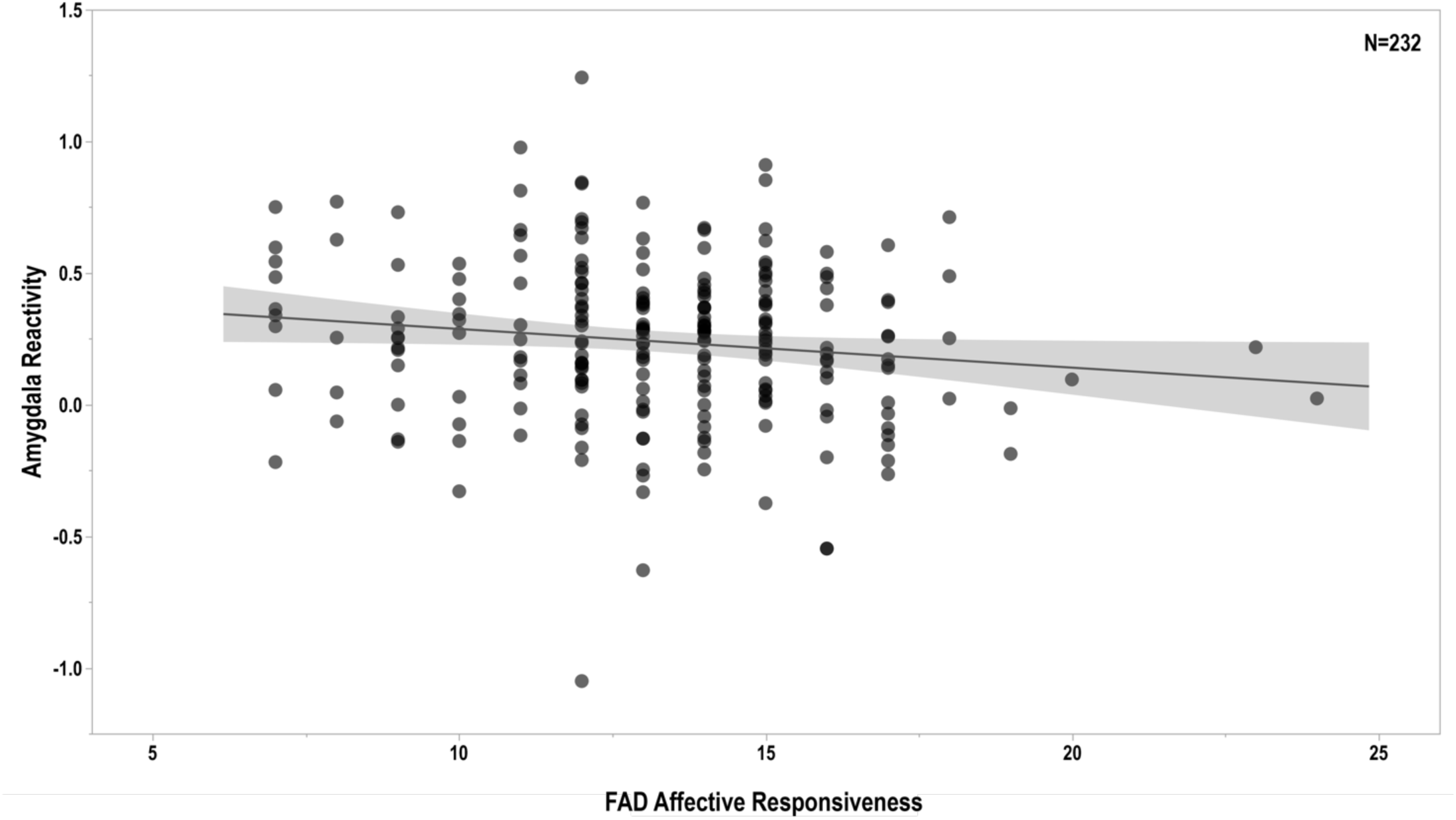
Family Functioning and Amygdala Reactivity. FAD affective responsiveness scores are negatively correlated with left amygdala reactivity to interpersonal threat as indexed by angry facial expressions [*F*(1, 221)=4.305, *p*=0.039]. Of note, higher scores on the FAD reflect poorer functioning, while lower scores reflect better functioning.

Importantly, planned post hoc analyses revealed there were no significant associations between amygdala reactivity to angry facial expressions and FAD general functioning scores [left: *F*(1, 221)=0.611, *p*=0.435; right: *F*(1, 221)=0.025, *p*=0.874], or between amygdala reactivity to fearful facial expressions and either general functioning [left: *F*(1, 221)=0.037, *p*=0.848; right: *F*(1, 221)=0.567, *p*=0.452] or affective responsiveness [left: *F*(1, 221)=0.011, *p*=0.915; right: *F*(1, 221)=0.493, *p*=0.483]. Lastly, regression analyses revealed a significant interaction of recent life stress and FAD affective responsiveness [*F*(1, 213)=4.581, *p*=0.033], such that the association between affective responsiveness and amygdala reactivity to angry facial expressions was moderated by the experience of recent life stress (Figure 3). Specifically, the association between better affective responsiveness and higher left amygdala reactivity to angry facial expressions was significant for participants reporting lower [*r*=-0.244, *p*=0.014] but not higher recent life stress [*r*=-0.052, *p*=0.614].

**Figure 3.**
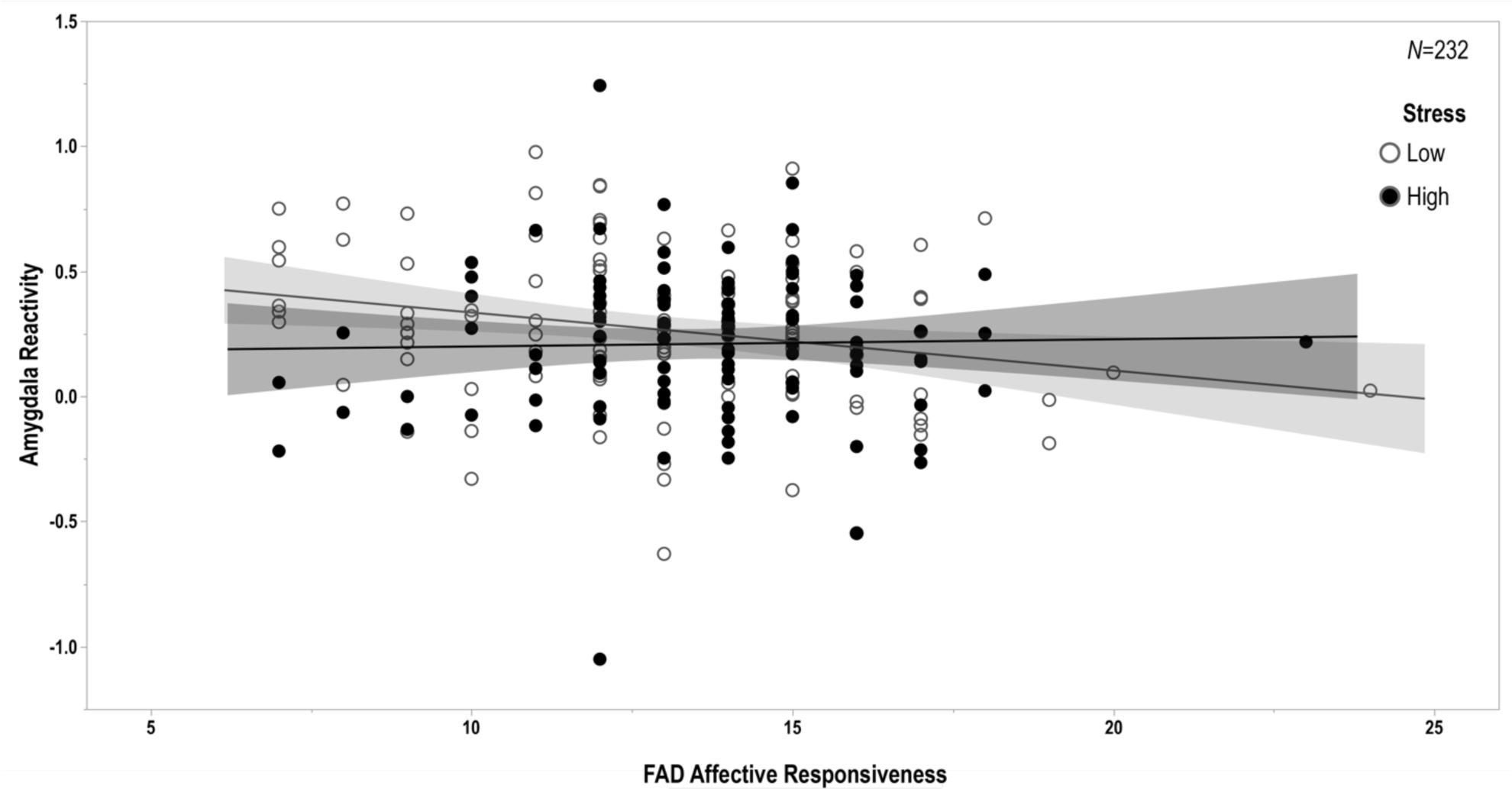
Family Functioning, Recent Stress, and Amygdala Reactivity. The negative correlation between FAD affective responsiveness and amygdala reactivity to interpersonal threat was significant only for participants reporting relatively low levels of recent stressful life events as assessed by the SLES [*F*(1, 213)=4.581, *p*=0.033; low stress: *r*=-0.244, *p*=0.014; high stress: *r*=-0.052, *p*=0.614]. Of note, higher scores on the FAD reflect poorer functioning, while lower scores reflect better functioning.

## Discussion

Our findings extend the literature on the impact of caregiving extremes on behaviorally-relevant brain function by demonstrating a significant association between normative variability in family functioning and threat-related amygdala reactivity. Specifically, we find that even subtle differences in caregiving within the family map onto variability in amygdala reactivity to explicit signals of interpersonal threat (i.e., angry facial expressions) but not implicit signals of broad environmental threat (i.e., fearful facial expressions). Contrary to our hypothesis, however, better familial affective responsiveness was associated with increased amygdala reactivity to interpersonal threat. This association was robust to the potential influence of participant sex, age, early life stress, contemporaneous symptoms of depression and anxiety, and broad familial risk for depression. Moreover, this association was moderated by the experience of recent stressful life events wherein better affective responsiveness was associated with higher amygdala reactivity in participants reporting low but not high recent stress.

Our paradoxical associations are particularly interesting when conceptualizing family functioning as an interpersonal process. For example, in families in which emotions are not appropriately processed, children may frequently experience interpersonal conflict including anger between family members and thus may be desensitized to interpersonal threat cues, as evidenced by lower amygdala reactivity to angry facial expressions. Alternatively, it is possible that children raised by parents who more appropriately experience, express, and model their emotions are sheltered from interpersonal conflict, making them more sensitive to interpersonal threat cues and thus more likely to exhibit amygdala hyper-reactivity to angry facial expressions. Consistent with this speculation, our findings revealed that the association between familial affective responsiveness and amygdala reactivity to angry expressions was specific to participants who reported having experienced relatively less recent life stress. Thus, adolescents who are raised in emotionally expressive, supportive family environments and who experience few external stressors may be more likely to exhibit neural hypersensitivity to interpersonal threat perhaps reflecting their lack of exposure to conflict.

Importantly, there is support for our observed paradoxical findings from both basic biobehavioral and clinical developmental research. For example, work by Fanselow & Tighe (1988) demonstrates that the length of time between aversive shocks has a greater impact on freezing behavior in rats than does the frequency of shocks. Specifically, unpredictable, infrequent shocks result in stronger freezing behavior than regular, frequent shocks. This pattern of potentiated aversive learning, which is mediated by amygdala circuitry, is more generally consistent with the increased effectiveness of spaced versus massed learning (Bouton, 2007). Thus, the observed amygdala hyper-reactivity to interpersonal threat cues in adolescents from families with better affective responsiveness who experience lower recent stress may reflect heightened associative learning following less-frequent, more unpredictable experiences of threat or conflict.

In parallel to this basic research, developmental clinical psychology studies have observed patterns between family functioning and threat-related biobehavioral processes similar to those we observe. For example, Andreotti et al. found that only young adults who reported lower levels of family conflict in early life exhibited increased cortisol reactivity to an acute laboratory stressor that subsequently predicted subliminal attentional bias to threat cues (Andreotti et al., 2015). Notably, the amygdala plays a significant role in both processing subliminal threat signals and triggering cortisol release via activation of the hypothalamic-pituitary-adrenal axis (Whalen & Phelps, 2009). More generally, parental overprotection, which may be characterized by either more restrictive and controlling (Parker, 1983) or indulgent (Green & Solnit, 1964) parenting has been associated with later psychopathology including depression and anxiety disorders (Thomasgard & Metz, 1993). Consistent with these patterns, amygdala hyper-reactivity is a core pathophysiological observation in these disorders (Etkin & Wagner, 2007) and has emerged as a biomarker of future risk (Admon et al., 2009; Mattson et al, 2016; McLaughlin et al., 2014; Swartz et al., 2014). It is interesting to further speculate that our observed associations between better familial affective responsiveness, less stress, and higher amygdala reactivity hints at a mechanism through which parental overprotection may manifest as psychosocial dysfunction.

To our knowledge, this is one of very few studies investigating the neural correlates of normative variability in caregiving in a non-clinical sample of adolescents. In addition to our current findings, Romund et al. (2016) have reported that increased maternal warmth and support are correlated with lower amygdala reactivity to fearful facial expressions amongst a healthy sample of adolescents, although they did not find an association with reactivity to angry facial expressions. The divergence of our associations with that of Romund et al. may reflect differences between broader, non-specific familial dynamics and maternal-specific behaviors, explicit modeling of recent stress, as well as differences in the task-based amygdala reactivity elicited. Of note, unlike our conservative approach using unbiased amygdala reactivity values originating from our main effects of expression contrasts in our analyses of a much larger sample (232 vs. 83), Romund et al. did not find significant main effects of angry or fearful facial expressions on amygdala reactivity. Rather, they conducted more permissive correlation analyses identifying specific voxels within the amygdala that exhibited BOLD values correlated with their measures of maternal caregiving.

In another recent study, Thijssen et al. (2017) reported that insensitive parenting was associated with a more mature pattern of intrinsic connectivity between the amygdala and medial prefrontal cortex, which together function to integrate and regulate emotional experiences, in healthy children. And in an earlier study, Taylor et al. (2006) reported that lower family stress during childhood was associated with lower correlated activity between the amygdala and regulatory regions of the ventrolateral prefrontal cortex during affective labeling of angry and fearful facial expressions in adulthood. Despite these differences, our findings and those of these prior studies suggest that normative variability in caregiving and family functioning is associated with alterations in threat-related brain function similar to those that have been reported for caregiving extremes including trauma, abuse, and neglect.

As exaggerated threat-related amygdala reactivity has been linked to a range of clinical outcomes, particularly following stressful life events (Swartz et al., 2015), next steps could include probing the relationship between family functioning, clinical outcomes following stressors, and amygdala function. Our initial evidence linking normative variability in caregiving with threat-related brain function can now serve as a starting point for pursuing longitudinal data analyses to investigate potential trajectories of amygdala reactivity and family functioning over time. Future research can further explore this question using whole-brain analyses, as well as examining functional connectivity between the amygdala and prefrontal cortex during the explicit regulation of negative emotions to gauge the impact of family functioning on top-down control processes.

Our study, of course, is not without limitations that can be addressed in future research. First, we relied on self-reported measures of caregiving using a single instrument. Similarly, we assessed symptoms of anxiety and depression using self-report inventories. Thus, our findings may be subject to reporting bias and are likely more representative of the perception of events rather than of objective events. Second, hemispheric lateralization of significant associations between caregiving and amygdala reactivity was not hypothesized and therefore awaits replication before further useful speculation. Third, we did not have an explicit measure of parental overprotection. Thus, our speculation regarding possible links between affective responsiveness and this aspect of family functioning previously linked with sensitization and maladaptive responses to stress outcomes requires explicit testing. Fourth, we only evaluated amygdala reactivity to threat-related facial expressions. It is possible that normative variability in affective responsiveness may also map onto individual differences in amygdala reactivity to reward-related social signals, namely happy facial expressions, or emotionally neutral facial expressions, and that these associations may reflect additional pathways of risk or resilience to psychopathology. Lastly, TAOS is not a population representative sample but rather selected based on familial risk for depression. Future studies in diverse samples are necessary to evaluate the extent to which our current findings are present more broadly and, consequently, useful for informing risk for psychopathology more generally.

Despite these limitations, our findings suggest that better familial affective responsiveness is associated with a paradoxical increase in amygdala reactivity to explicit signals of interpersonal threat (i.e., angry facial expressions), especially in adolescents reporting lower recent stressful life events. Better familial affective responsiveness may exacerbate neural reactivity to interpersonal threat—particularly when coupled with fewer external stressors—by limiting opportunities for the expression and subsequent self-regulation of negative emotions. While our findings extend a rich literature on the effects of extremes in caregiving on emotional brain function to more subtle, normative variability they also indicate that a more careful consideration of putatively protective factors in the expression of risk-related brain function may be necessary in the absence of caregiving extremes.

## Materials & Methods

### Participants

Study participants were recruited into two groups: “high risk” wherein a first- and second-degree relative has a history of MDD, or “low risk” wherein no first-degree and minimal second-degree relatives (<20%) have such a history. Low risk participants further could not meet criteria for any psychiatric disorder or substance use disorder; however, a diagnosis of an anxiety disorder was permitted in the high risk participants as early-onset anxiety is frequently a precursor to depression (Goodwin et al., 2004; McLaughlin & King, 2015). Mental health in all participants was assessed through structured clinical interviews with the adolescent and parent separately using the Kiddie Schedule for affective Disorders and Schizophrenia for School-Age Children Present and Lifetime Version (Kaufman et al., 1997). All procedures were approved by the Institutional Review Board at the University of Texas Health Sciences Center San Antonio, and participants and their parent provided written informed assent/consent before participation. Additional recruitment details have been described in detail elsewhere (Bogdan et al., 2012; Swartz et al., 2017; White et al., 2012).

Of the 331 participants completing the parent study, Teen Alcohol Outcomes Study, data for our present analyses were available for 232 participants (120 high risk, 112 low risk). Of these 331 participants, data were excluded for 29 due to problems with the scan or raw data, (e.g., ending the scan early, poor coverage of the amygdala, gross anatomical abnormalities, scanner artifacts). Data from another 7 high risk participants were excluded because they had a diagnosis of MDD before the scan, and from another 5 low risk participants because they had an anxiety disorder before the scan. Functional MRI data from the remaining 290 participants underwent pre-processing and, subsequently, another 53 participants were excluded based on quality control criteria for the processed data (see quality control procedures section below for further details). Lastly, data from another 5 participants were excluded for poor accuracy on the fMRI task. The final analysis sample of 232 participants was 50.4% female and approximately 58% Caucasian, 26% Hispanic, 5% African American, and 12% Indian, Asian, Pacific Islander, or multiple races. There was no difference in sex ratio (χ^2^(1)=0.018, *p*=0.892) or the distribution of race/ethnicity (χ^2^(4)=2.413, *p*=0.660) between the high risk and low risk groups.

### Behavioral Measures

Family functioning was assessed using the Family Assessment Device (FAD), a 60-item scale completed by both child- and parent, which yields 7 scales of family functioning. Family members rate how each item describes their family by selecting ‘strongly agree’, ‘agree’, ‘disagree’, or ‘strongly disagree’. Scores per item range from 1 to 4 with 1 reflecting healthy functioning and 4 reflecting unhealthy functioning (Epstein et al., 1983; Stein et al., 2000). Given the literature on the impact of warmth and care provided by caregivers, we probed this feature of caregiving on amygdala reactivity specifically using the “affective responsiveness” subscale, which measures the appropriate experience and expression over a range of situations, with items such as “We are reluctant to show our affection for each other” and “We express tenderness.” We addressed the possible unique impact of this caregiving feature on amygdala reactivity by also examining correlations with the “general functioning” subscale, which measures the overall level of family functioning with items such as “In times of crisis we can turn to each other for support” and “We don’t get along well together.” We utilized child- and not parent-reported scores from these two subscales because we expected that a child’s perception of their familial environment would be more impactful than the parent’s perception, and we wanted to better approximate scores from the Parental Bonding Instrument, which is a more commonly used inventory of normative caregiving that is child-reported, but unavailable in the present study. The FAD has been extensively used in a variety of research contexts and has shown good test-retest and concurrent reliability as well as good internal consistency and validity (Epstein et al., 1983).

Depression symptoms were assessed using the Mood and Feelings Questionnaire (MFQ), a widely-used measure in children and adolescents (Messer et al., 1995). We used the child-reported short (13 item) summary score in our analyses. Anxiety symptoms were assessed using the Screen for Child Anxiety Related Disorders (SCARED), a widely-used measure of anxiety symptoms in children and adolescents (Birmaher et al., 1999). We use the child-reported SCARED in our analyses.

Early life stress was assessed using the Childhood Trauma Questionnaire (CTQ), a widely-used measure of childhood trauma and early life stress (Bernstein et al., 2003). Recent stress was measured using the Stressful Life Events Schedule (SLES), an interview designed to assess for life stressors relevant to children and adolescents (Williamson et al., 2003).

### Functional MRI

Participants performed an emotional face matching task that has been shown to consistently elicit robust amygdala activity in numerous studies of adults and adolescents (Bogdan et al., 2012; Swartz et al., 2017; White et al., 2012). Task blocks consisted of matching angry and fearful facials expressions while control blocks consisted of matching geometric shapes (Bogdan et al., 2012; Swartz et al., 2015; Swartz et al., 2017; White et al., 2012). Further task details are reported in the supplement.

Analyses were conducted using the general linear model of SPM8 (http://www.fil.ion.ucl.ac.uk/spm/software/spm8/). Following preprocessing, linear contrasts employing canonical hemodynamic response functions were used to estimate main effects of expression for each individual. Individual contrast images were then entered in second-level random effects models to determine mean condition-specific regional responses using one-sample t-tests. We extracted parameter estimates from functional clusters within anatomically defined amygdala regions of interest (Automated Anatomical Labeling atlas) at *p*<0.05 family wise error (FWE) corrected across the search volumes for the contrast of angry blocks > control blocks and fearful blocks > control blocks (Swartz et al., 2015). Further quality assurance and extraction details are reported in the supplement.

### Statistical Analyses

Mean individual contrast-related BOLD activation values from functional clusters were entered into second-level analyses in SPSS, version 24 (IBM, Armonk, N.Y.). First, we ran our primary linear regression of the FAD affective responsiveness subscale with extracted BOLD values for the contrast of angry expressions greater than shapes. We then conducted analyses to confirm that the significant association between familial affective responsiveness and amygdala reactivity was not driven by early life adversity or concurrent depressive and anxiety symptoms by duplicating regression analyses from step one, this time controlling for CTQ, MFQ and SCARED scores in addition to our other covariates. To probe the specificity of this association, we conducted post hoc linear regressions of the affective responsiveness subscale with extracted BOLD values for the contrast of fearful expressions greater than shapes, as well as linear regressions of the FAD general functioning subscale with extracted BOLD values for each expression-specific contrast. Lastly, we conducted linear regressions to investigate potential direct effects of recent stress (SLES) and interactions of SLES with FAD. We then conducted post hoc partial correlation analyses within the high and low stress groups to test for significance of the simple slopes. In all analyses, we controlled for participant age, sex, and risk group.

## Acknowledgments

We thank the Teen Alcohol Outcomes Study participants as well as the staff of the Laboratory of NeuroGenetics and the Williamson Laboratory. The main funding for this study was supported by the R01AA016274 (Williamson). The Laboratory of NeuroGenetics received support from Duke University as well as US-National Institutes of Health Grants R01DA033369, R01DA03157, and R01AG049789.

